# Comparing the advantages and disadvantages of physics-based and neural network-based modelling for predicting cycling power

**DOI:** 10.1101/2023.08.08.552425

**Authors:** Patrick Mayerhofer, Ivan Bajić, J. Maxwell Donelan

## Abstract

Models of physical phenomena can be developed using two distinct approaches: using expert knowledge of the underlying physical principles, or using experimental data to train a neural network. Here, our aim was to better understand the advantages and disadvantages of these two approaches. We chose to model cycling power because the physical principles are already well understood. Nine participants followed changes in cycling cadence transmitted through a metronome via earphones and we measured their cadence and power. We then developed and trained a physics-based model and a simple neural network model, where both models had cadence, derivative of cadence, and gear ratio as input, and power as output. We found no significant differences in the prediction performance between the models. The advantages of the neural network model were that, for similar performance, it did not require an understanding of the underlying principles of cycling nor did it require measurements of fixed parameters such as system weight or wheel size. These same features also give the physics-based model the advantage of interpretability, which can be important when scientists want to better understand the process being modelled.

## Introduction

Models are useful. Using models, scientists can better understand and prevent injuries (Aguiar et al., 2021; Ayala et al., 2019; Finch, 2006; Hulme et al., 2019; Souza & Gottfried, 2013; Spörri et al., 2017). Models can give insights into how muscles work (González-Izal et al., 2012; Hill, 1938; Rahemi et al., 2014; Schellenberg et al., 2015; Ting et al., 2012) and help define human limits to develop anti-doping tools (Faiss et al., 2019; Montagna & Hopker, 2018; Puchowicz et al., 2018). They can also simulate variables that are hard or expensive to measure. In cycling, for example, devices use models to estimate the mechanical power output of riders. This is useful for riders who do not own equipment that can measure mechanical power directly.

Developing these models from first principles can be difficult. Scientists design models to describe the real-world relationships that transform independent variables into the dependent variables. One option to model this input-output relationship is to understand and apply the principles underlying a particular process. For example, developing a physics-based model for cycling that can predict mechanical power (the output variable) requires first identifying the input variables that affect the power—such as speed or drag forces—and then identifying the parameters and how they are combined with the input variables to predict the output variable (Martin et al., 1998). These parameters can be identified through measurements, or from data (Dahmen et al., 2012). Using this physics-based approach to develop models has at least two major challenges. First, this process can require a detailed understanding of the principles underlying a process, which may be unknown or complex. Second, real-world measurements introduce inaccuracies, which stem from equipment measurement errors, and the fact that nothing can be measured perfectly. Inaccurately measured parameters can subsequently reduce the performance of the overall model. For example, to predict cycling speed from cadence a scientist would have to first understand the underlying principles of how the angular velocity of the pedal translates to the linear velocity of the wheel, and second to measure bike parameters such as the wheel radius.

Data-driven neural networks can assist the process of developing models. Neural networks are named as such because they loosely imitate the human brain. Mathematically, they consist of a set of functions that can be trained with data to recognize patterns in complex data sets. Because neural networks can learn from data, there are many applications for which they can be used. Modelling the input-output relationship of processes with neural networks still requires knowledge about the input variables. But, given enough data, neural networks can approximate a wide variety of input-output relationships without explicitly having to measure many relevant fixed parameters, or understand the principles underlying a process, ameliorating the two challenges identified above (Hornik et al., 1989). For example, to predict cycling speed from cadence a neural network could learn the relationship between the cadence and the cycling speed without understanding the principles that relate speed and drag to power, and without requiring measurements of the bike parameters such as the wheel radius.

In order to better understand the advantages and disadvantages of developing models using a first principles or data-driven approach, here we compare a physics-based model for predicting mechanical power in cycling with a predictive model developed using neural networks. We chose cycling for two reasons. First, the underlying principles upon which to build a physics-based model are well-understood (Debraux et al., 2011; Fitton & Symons, 2018; Maier et al., 2017; Martin et al., 1998). Second, there is both scientific and commercial interest in accurately predicting mechanical power during cycling. Scientifically, it can help to better simulate racing strategies (Dahmen et al., 2012; Fitton & Symons, 2018; Gordon, 2005; Wolf et al., 2016). Commercially, this knowledge can lead to products that help athletes to indirectly measure their power without the necessity of expensive power metres. To accomplish our goal, we first built a microcontroller-operated system that provides the cyclists with metronome-indicated changes in cadence and measures the power output. We used this system to measure cyclists’ power output during two trials of cycling on a flat running track with two different gear ratios while following step changes in commanded cadence. Using this data, we developed and parameterized a physics-based model and a neural network model that best fit the simulated power to the measured power. We then compared how accurately these two models predicted the measured mechanical power, and evaluated the advantages and disadvantages of each approach.

## Methods

### Data Collection

We tested nine participants in this experiment (3 females and 6 males; body mass: 72.2 ± 9.4 kg; height: 177.2 ± 10.5 cm; age: 28 ± 5 years; mean ± std). The Office of Research Ethics at Simon Fraser University approved the study (#20180650). All participants provided written and verbal informed consent before participating in our study.

During the experiment, participants cycled on a 400 m running track. All participants used the same bike (Specialized Tricross Comp Size 52, Specialized Bicycle Components, Inc.) and adjusted the seat height to their own preference. During cycling, they carried a backpack with a microcontroller (Teensy 3.1, Pjrccom Llc.). The microcontroller measured torque in the pedal crank arm *τ*_*p*_ continuously from an SRM power metre (Dura-Ace, SRM GmbH). Twice per crank arm revolution (every half pedal stroke), the microcontroller measured the crank arm angular velocity *ω*_*p*_ using a reed switch, and calculated the mechanical power *P*_*P*_ by multiplying the time-averaged torque with the angular velocity:

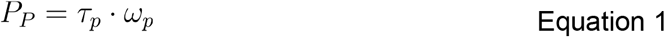

As a warm-up and to familiarise participants with the bike, we first instructed them to cycle at a comfortable speed for 5-10 minutes. During this familiarisation period, participants chose their preferred rear gear (16.5 ± 0.5 teeth), while we kept the front gear fixed (39 teeth). We measured their preferred cadence (67 ± 12 rpm) as the average cadence during a 30 second period towards the end of the familiarisation period. Next, participants completed an 18 minute trial with the rear gear being one gear over their preferred rear gear. This was followed by a second 18 minute trial with the rear gear one gear under their preferred rear gear. We instructed participants to keep their body position (i.e.: high vs. low handlebar position) the same throughout the experiment to keep their frontal area, which affects the drag, relatively constant. A metronome, controlled by the microcontroller and communicated to the participant through earphones, commanded step changes of ±5%, ±10%, and ±20% of the participant’s preferred cadence, centred about the preferred cadence (Figure 1). We instructed participants to match the metronome beat as accurately as possible with their cadence. Step changes occurred every 60 s and participants could rest for ∼10 mins between the two trials.

**Figure 1:**
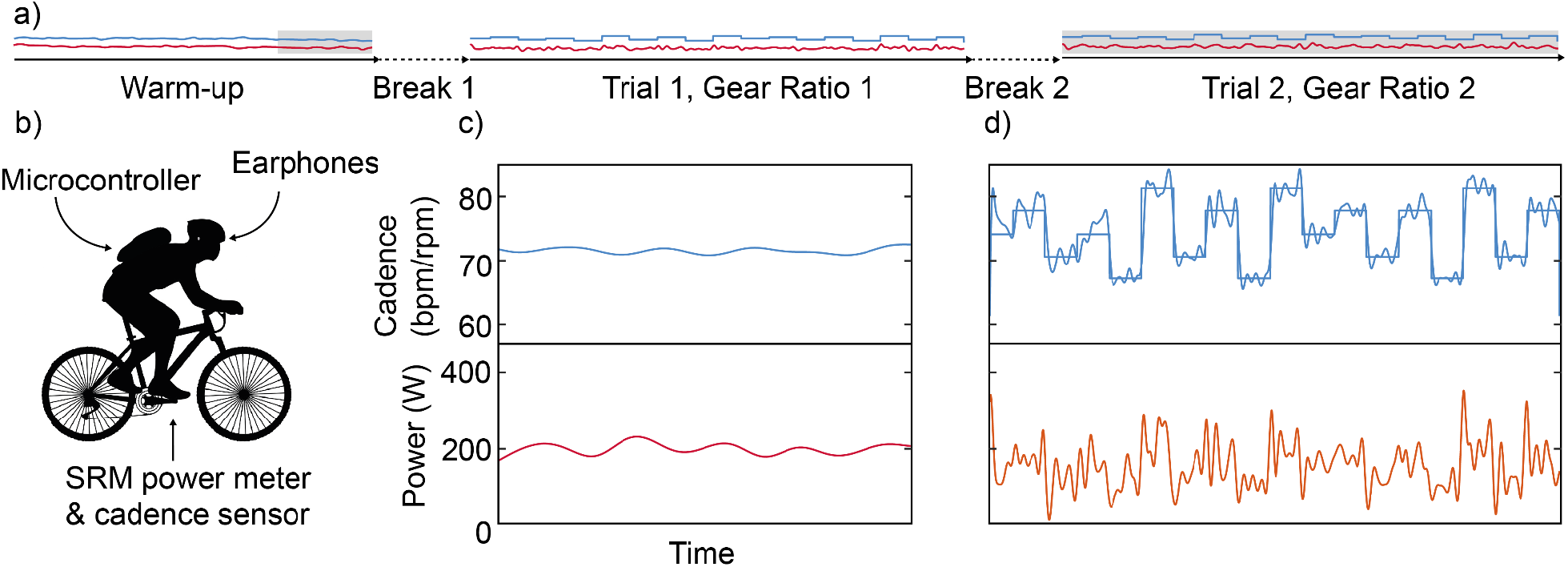
The experimental setup. a) illustrates the timeline of the experiment with the Warm-up, during which we evaluated the participant’s preferred cadence (grey box), Break 1, Trial 1 with Gear Ratio 1, Break 2, and Trial 2, with Gear Ratio 2. b) illustrates the participant with the equipment. c) magnifies the data in the grey box of the warm-up. d) magnifies the data in the grey box of trial 2. Metronome cadence is measured in beats per minute (bpm), cycling cadence is measured in revolutions per minute (rpm), and power is measured in Watts.

### Development of the Physics-based Model

To derive an expression for the mechanical output power of the cyclist as a function of the input cadence and gear ratio of the cyclist we model the system as horizontal forces acting on a point mass (*m*). One force is applied by the bike’s rear wheel (F_*rw*_), and a counteracting force is applied by air resistance (F_*drag*_). Using Newton’s second law to describe the cyclist’s motion yields:

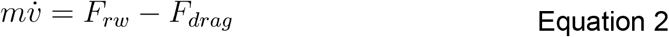

Where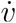 is the rate of change of the cyclist’s speed. *F*_*draw*_ is dependent on the squared speed (*υ*^2^), the air density (*ρ*), the frontal area of the cyclist (*A*), and the drag coefficient (*C*_*d*_) (Benson, 2021):

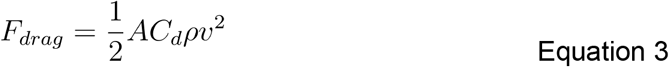

Due to multiple reasons, we replaced all but the squared velocity with one single variable which we call the drag number (*c*). First, we did not measure frontal area or air density. Second, our drag number is not only air drag but subsumes all other factors of drag, such as rolling resistance. Isolating *F*_*rw*_ in equation 2, and substituting the product of the squared speed and the drag number for *F*_*drag*_, yields:

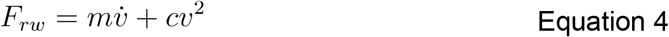

Using the forces and radii in Figure 2 to calculate the transformation of force from the pedal to the rear wheel yields:

**Figure 2:**
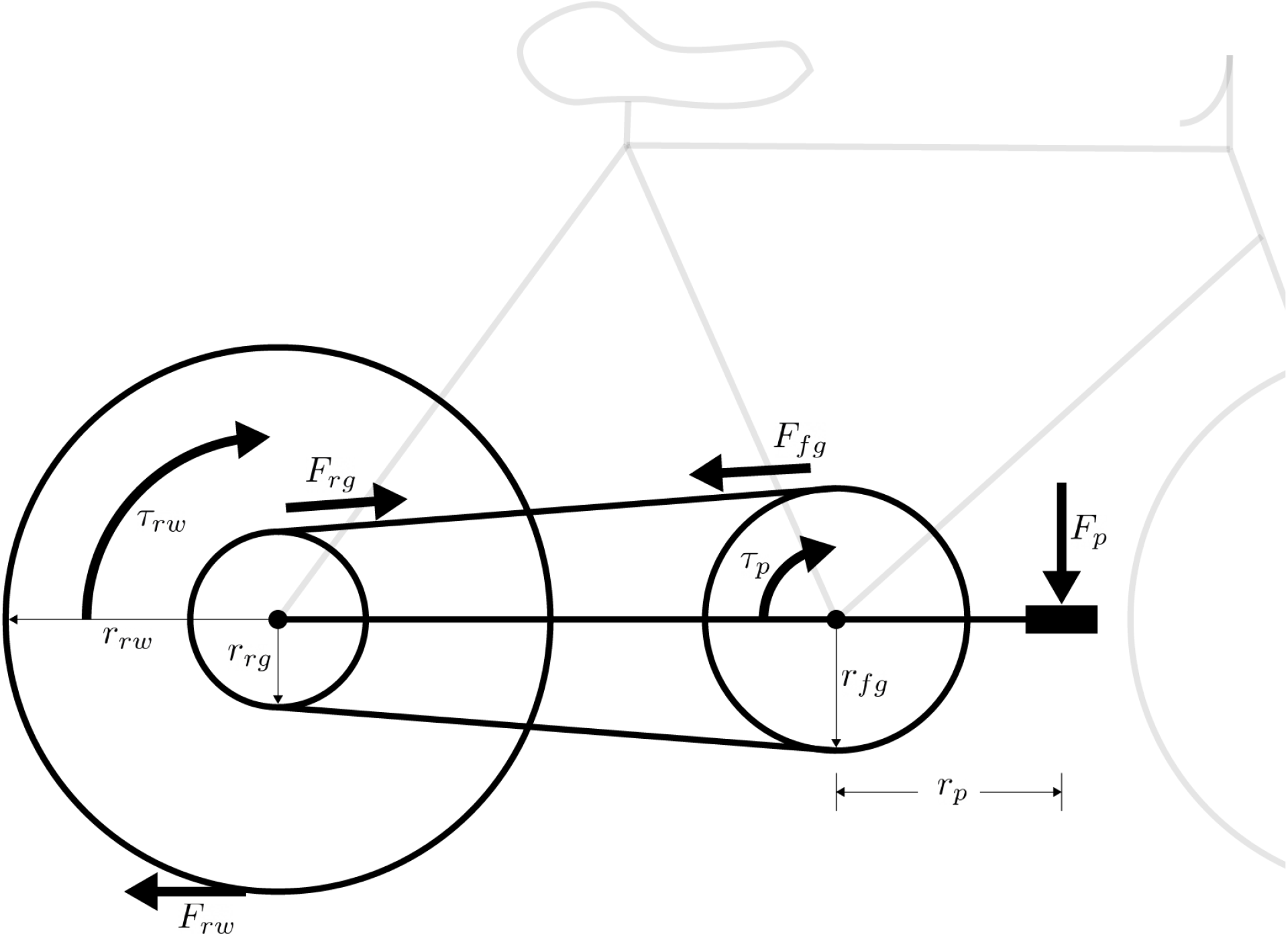
Illustrates the relevant forces (*F*), radii (*r*), and torques (*τ*) in the pedal, front gear, rear gear, and rear wheel.

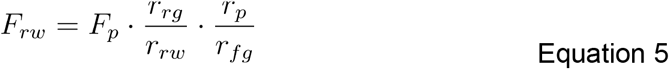

Replacing *F*_*p*_ with the ratio between the torque generated by the user on the crank arm (*τ*_*p*_) and the pedal length (*r*_*p*_) and the ratio between the front gear radius (*r*_*fg*_) and the rear gear radius (*r*_*rg*_) with the gear ratio (*GR*) yields:

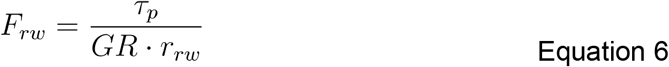

Substituting this expression for *F*_*rw*_ into equation 4 yields:

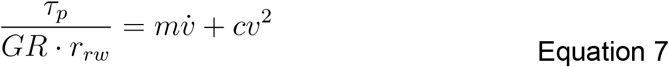

Multiplying both sides with the gear ratio, the rear wheel radius, and the pedal’s angular velocity yields the mechanical power applied by the cyclist (*P*_*p*_) on the left side of the equation:

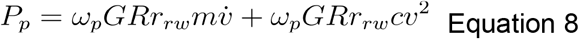

The angular velocity in the pedal is equal to the ratio between the angular velocity in the rear wheel and the gear ratio, and the angular velocity in the rear wheel equals the linear velocity of the cyclist and the radius of the rear wheel, yielding:

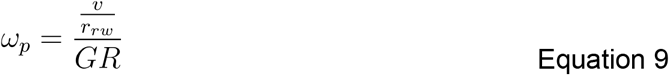

Substituting this expression for *ω*_*p*_ into equation 8 simplifies that equation to:

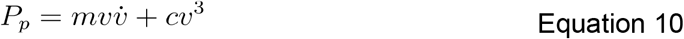

Expressing speed as a function of the measured gear ratio and the measured cadence (*f*) yields:

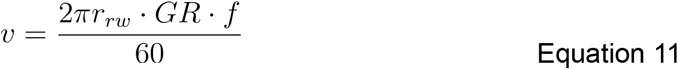

Substituting this expression for *v* into equation 10 and simplifying yields:

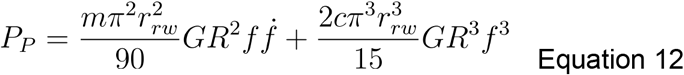

where 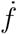 is the rate of change of the cadence. This equation expresses the output mechanical power of a cyclist as a function of the measured time-varying cadence and the experimentally-manipulated gear ratio. The only unknown and optimizable parameter is the drag number *c* — all other parameters in the equation can be measured (Table 1). To optimise for a drag number that best fit the predicted power to the measured power, we used a Levenberg-Marquardt optimization algorithm, implemented in Matlab’s nlinfit function (R2020a, The MathWorks, Inc.) (Seber & Wild, 1989).

**Table 1:** Values of all measurable parameters of the physics-based model.

| Parameters |  |
| --- | --- |
| $r_{rw}$ | 0.30m |
| $l$ | 0.17m |
| $m$ | 72.2±9.4kg (participant weight) + 13.0kg (bike and equipment weight) |
| $GR$ | 2.4±0.2 |

### Development of a Neural Network Model

When developing a neural network, there are many choices to make about the architecture of the network. These choices include the structure of the data that is input into the network, types of network layers, the number of layers, the number of nodes per layer, and the type of activation function applied to each layer’s output. While there are no clear rules to specify network architecture to maximise model performance on a given problem (Fiszelew et al., 2007; Hunter et al., 2012), there are certain architectures that have historically performed better on some problems than others. We used historical performance as well as pilot analyses to guide the following choices:

#### Data structure

To predict power for each half pedal stroke, we used input data from that half pedal stroke as well as the seven previous half pedal strokes. For all models, the input data at each half pedal stroke included the cadence and the derivative of cadence. For some models, the input data also included gear ratio (Figure 3). We chose time windows of eight half pedal strokes because longer windows required greater computational power and pilot analyses revealed good performance with our chosen window length.

**Figure 3:**
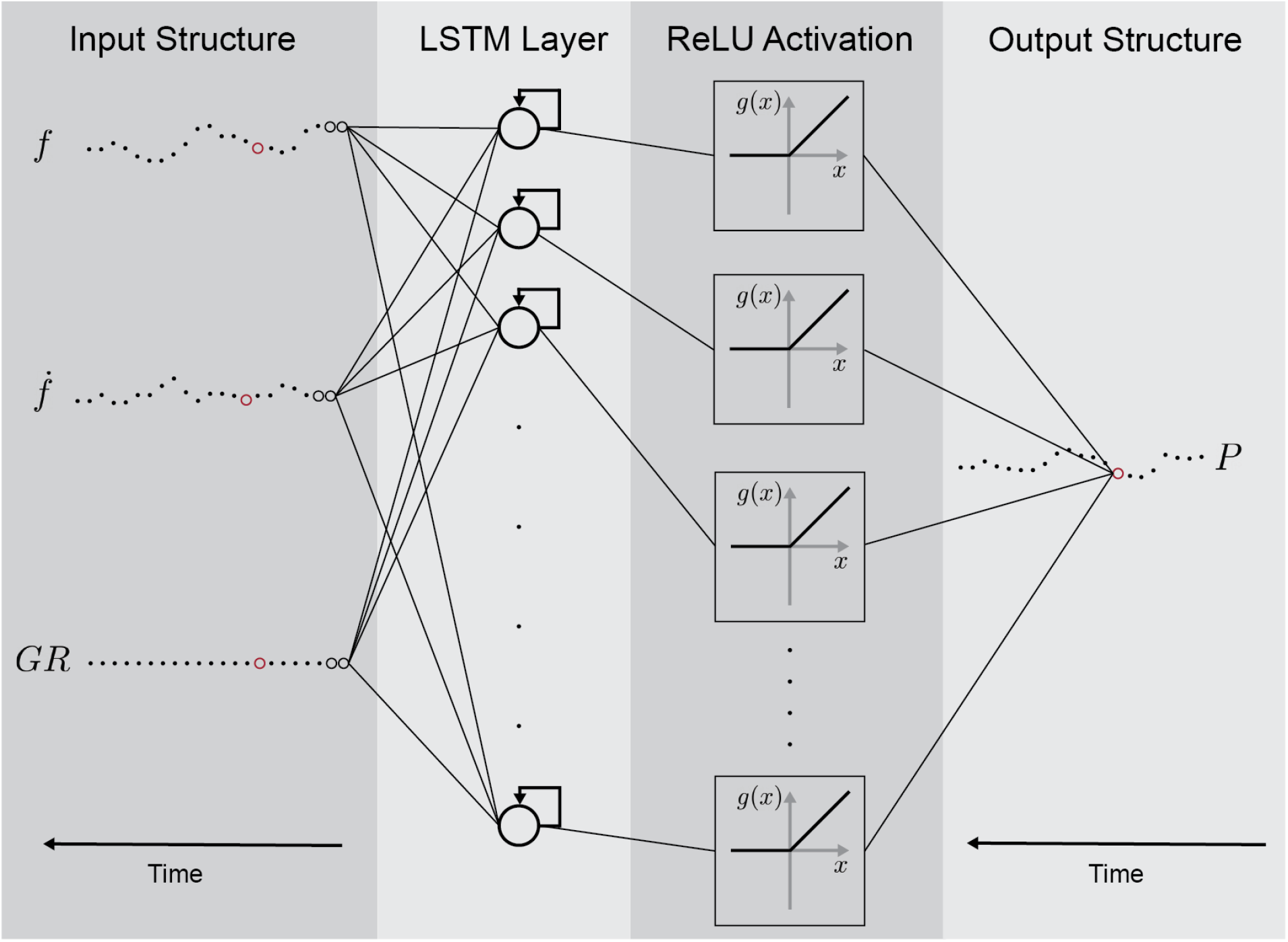
Illustration of a conceptual model of the neural network from input structure to output structure. Notice that for illustration purposes the data structure of the input and output are illustrated with the time evolving from right to left. To predict one output datapoint (power *P*) on the right (output structure), eight input time steps per input (cadence *f*, cadence derivative 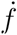, and gear ratio *GR*) are required. The red dots in the input structure illustrate the eighth and last input datapoint, and the red dot in the output structure illustrates the eighth output datapoint, which is the datapoint the neural network is predicting.

#### Types of layers

We chose to use recurrent layers which are comprised of nodes whose output can affect the next input to nodes of the same layer. They often perform better with temporal tasks—tasks where the data changes over time—because they can store information from past data. More specifically, we used long-short-term-memory layers, which can further improve the performance over other types of recurrent layers, by prioritising which information from past data to store (LeCun et al., 2015).

#### Number of layers

There are different advantages and disadvantages for both shallow neural networks (one hidden layer only) and deep neural networks (two or more hidden layers) (Bianchini & Scarselli, 2014; Kim & Gofman, 2018; Mhaskar et al., 2017). For simplicity, and because pilot analyses showed good performance, we chose to use only one hidden layer.

#### Number of nodes per layer

In pilot experiments, we found similar performance for a small number of layer nodes (8) when compared with greater numbers of nodes (16, 24,…, and 1024). For simplicity, we chose to use 8 nodes.

#### Activation function

Typically, each layer in a neural network is followed by an activation function, which transforms the output of each node in the layer and provides the network with non-linear modelling capabilities. Due to their widespread success in deep neural networks, we used a rectified linear unit (ReLU) as activation functions for the long-short-term-memory layer (Ramachandran et al., 2017; Sharma et al., 2017):

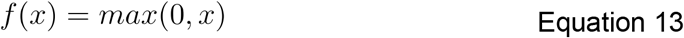

A ReLU activation function deactivates nodes with an output of smaller than 0, giving them the advantage of turning individual nodes on and off. We did not include additional activation functions for the output layer, because pilot analyses revealed better performance without output layer activation functions when compared with a ReLU activation function.

#### Data Analysis

To better determine the advantages and disadvantages of the physics-based and neural network models, we performed two analyses: a within-trial analysis and a within-participant analysis. First, we tested the prediction performance within each trial. Here the models’ aim was to learn from parts of the data within a trial and predict the rest of the data within the same trial. The neural network’s input was the cadence and the derivative of the cadence and did not require knowledge of the gear ratio, as it was a fixed parameter. Second, we tested the prediction performance within each participant. Here, the models’ aim was to learn from parts of both trials and predict the rest of the data within the same participant. Here, the neural network required knowledge of the gear ratio as an additional input, as it was a variable that was different between the two trials.

### We used normalised root mean square error, k-fold cross validation, and paired t-tests to compare model performance

To test the prediction performance, we calculated the normalised root mean square error, where we normalised the root mean square error by the mean of the measured data. Additionally, we also calculated the normalised mean error, where we normalised the mean error by the mean of the measured data. We split up each participant’s trial into three subsets, also called folds (Figure 4). To test the performance of the physics-based model and the neural network model in the within-trial experiment we trained the models with two of the subsets within a trial and tested the accuracy of predicting the power with the third, using both the normalised root mean square error and normalised mean error. For example, we would use fold 1a and fold 1b to train the model and fold 1c to test the prediction accuracy. Here, we did 3-fold cross validation: We used each of the three subsets as a test set once to get the prediction accuracy three times. To test the performance in the within-participant experiment, we trained the models with five of a participant’s subsets and tested the accuracy of predicting the power on the sixth. For example, we would use fold 1a, fold 1b, fold 1c, fold 2a, and fold 2b to train the model and fold 2c to test the prediction accuracy. Here we did 6-fold cross validation: We used each of the six subsets as a test set once to calculate the prediction accuracy six times. To compare overall performance, we averaged the normalised root mean square errors of each participant and compared the mean normalised root mean square error between the physics-based model and the neural network model with a paired t-test using a significance level of p < 0.05.

**Figure 4:**
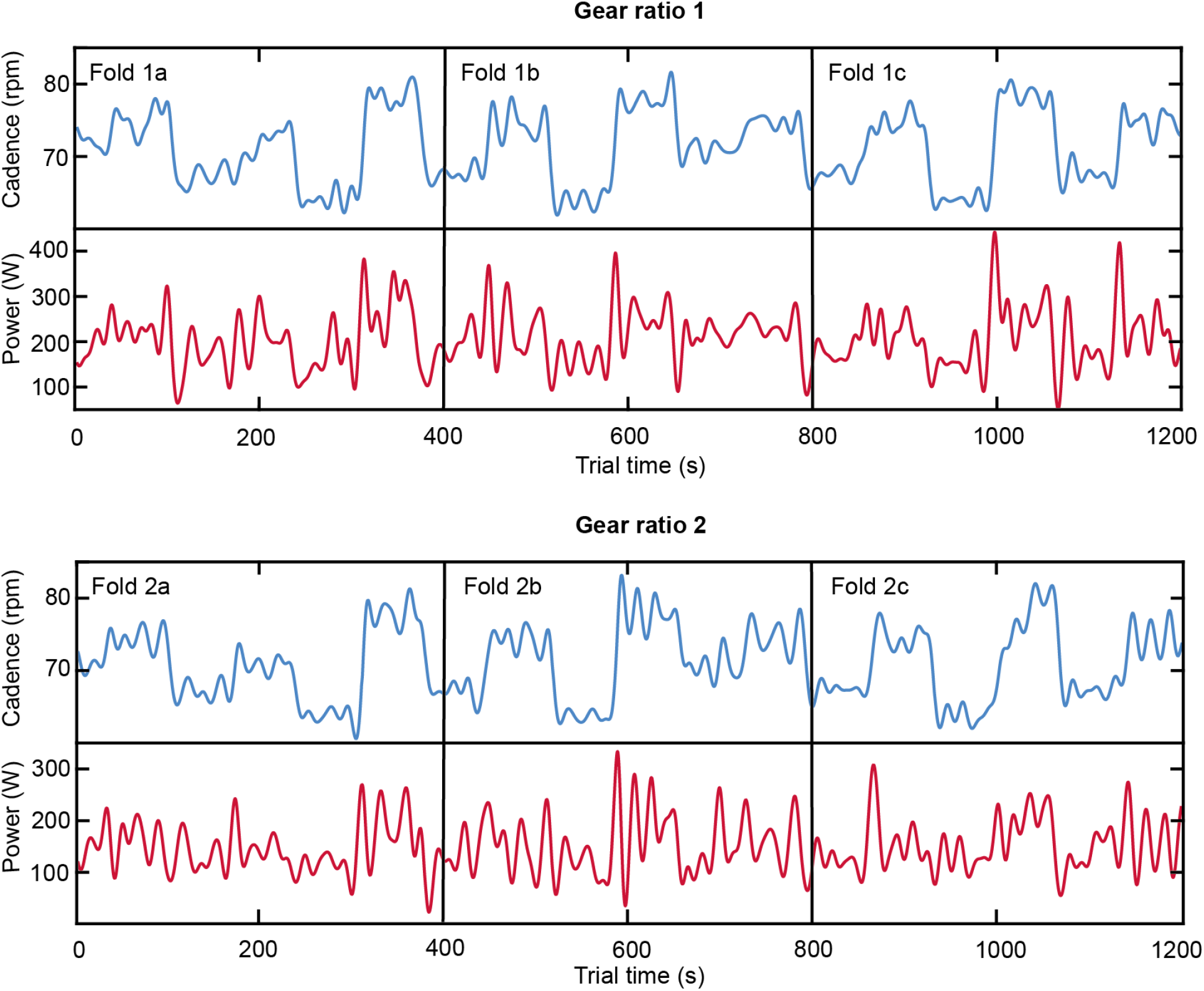
Illustrates the two trials with different gear ratios, and the subsets within each trial for the within-trial and within-participant analysis.

## Results

The physics-based model and the neural network model had similar predictive performance. The normalised mean error and normalised root mean square error for the within-trial analysis — in which the different gear ratio trials were kept separate when we trained and tested the model — of the physics-based model were 1.6%±1.1% and 19.6±5.1%, respectively (mean between participants ± standard deviation between participants; Figure 5). With this predictive performance, we expect a new participant with a measured average power output of 300 W to have a predicted average power that is ∼5 Watts (1.6%) above the actual average power. And for 95% of the half pedal strokes at 300 W, we expect the predicted power for this new participant to be within ∼116 Watts (19.6%*1.96). On average, the optimised drag number was 1.2±0.2. The normalised mean error and normalised root mean square error for the within-trial experiment of the neural network model were 1.2%±4.2% and 18.2±6.0%, respectively (Figure 5). The predictive performance between the two models was not significantly different (p = 0.34).

**Figure 5:**
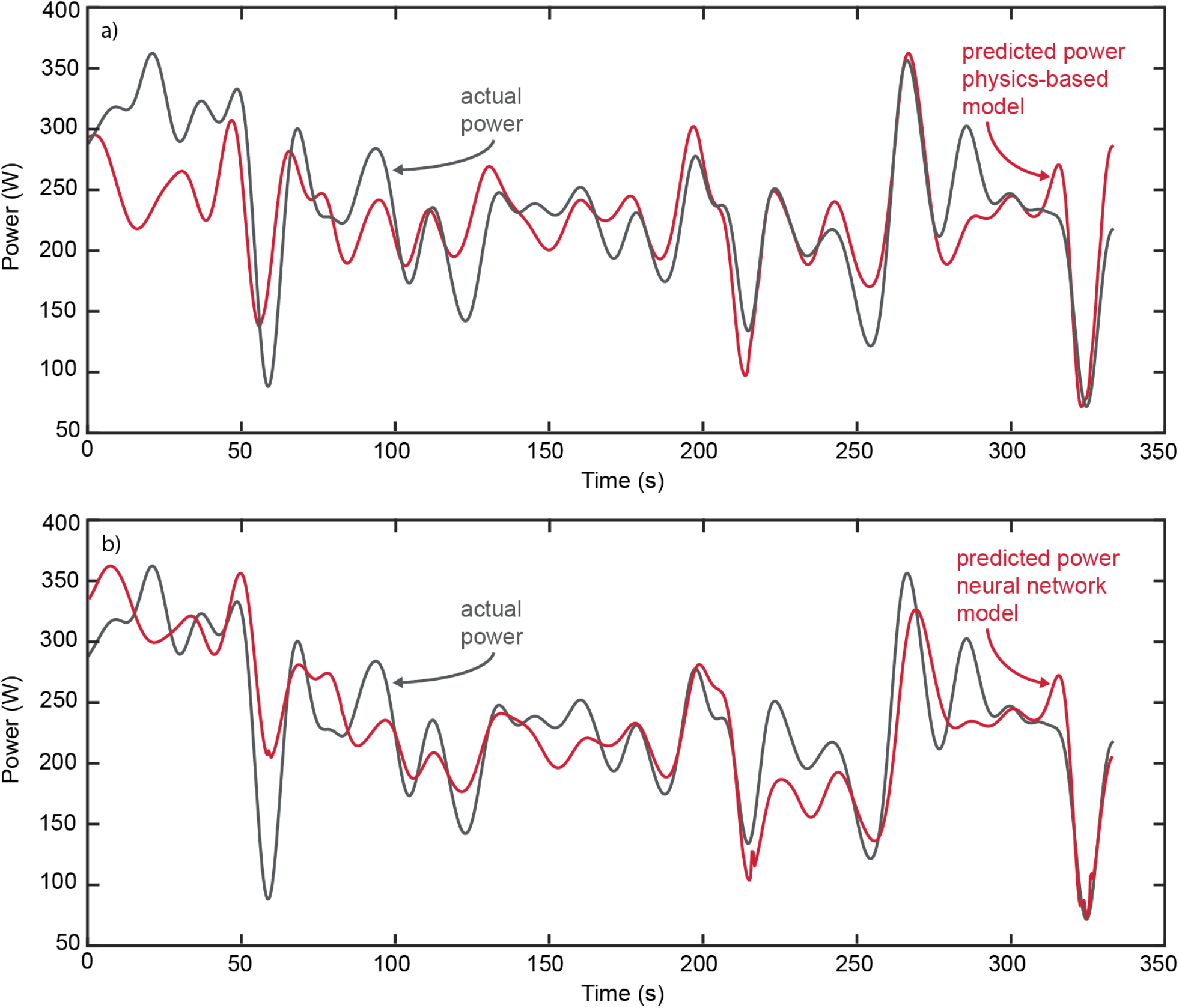
Representative prediction data for the within-trial analysis. These representative trials had similar RMSEs and normalised mean errors with the overall average.

The normalised mean error and normalised root mean square error for the within-participant analysis — in which we combined the different gear ratio trials when we trained and tested the model — of the physics-based model were 3.2% ± 1.8% and 20.9±5.1%, respectively (Figure 6). On average, the optimised drag coefficient was again 1.2 ± 0.2. The normalised mean error and normalised root mean square error for the within-participant experiment of the neural network model were 4.1%±10.9% and 25.4±5.6%, respectively (Figure 6). Again, the predictive performance between the two models was not significantly different (p = 0.12).

**Figure 6:**
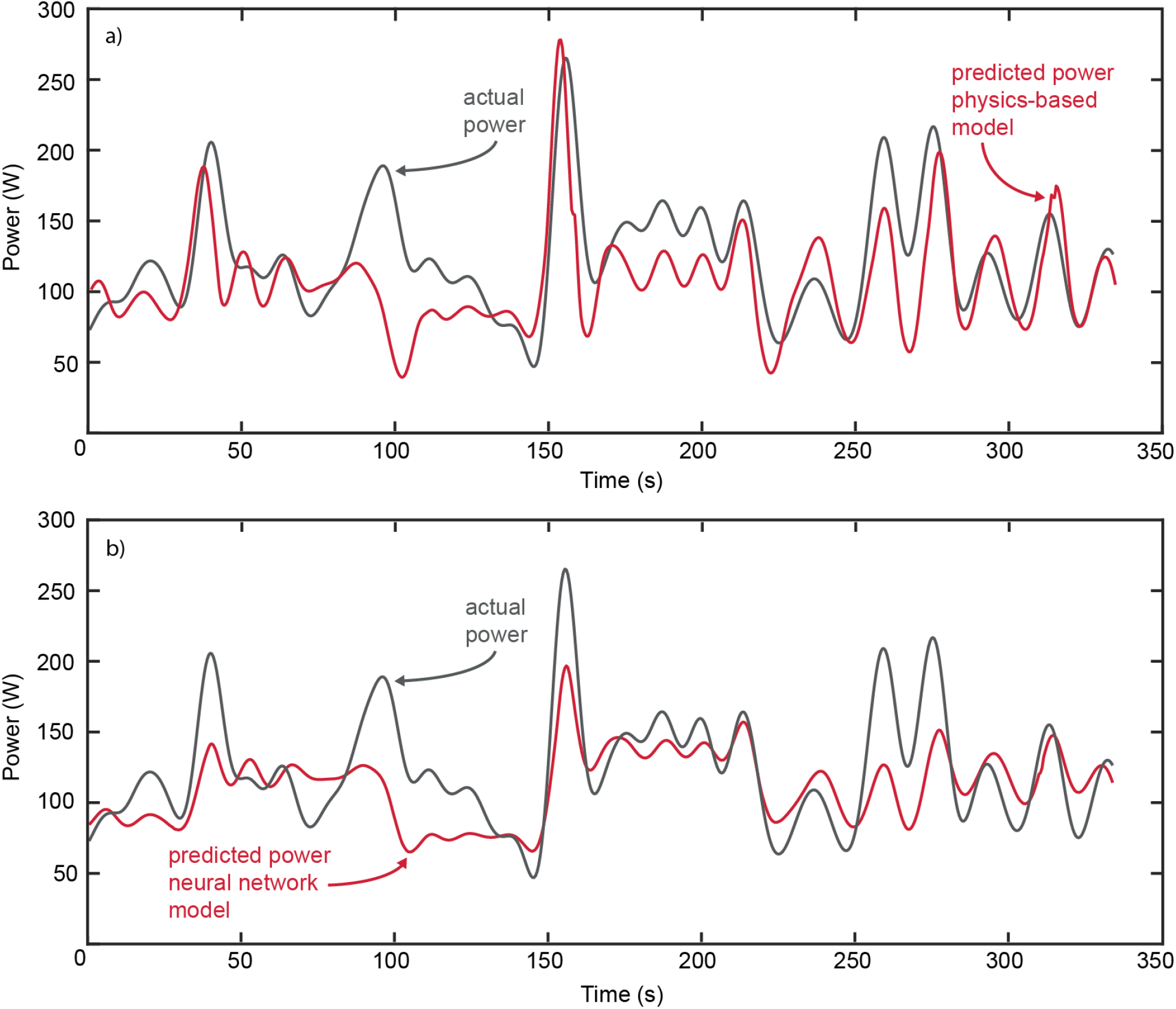
Representative prediction data for the within-participant analysis. These representative trials had similar RMSEs and normalised mean errors with the overall average.

## Discussion and Implications

Here we developed and compared a physics-based model and a neural network model in predicting cycling power from changes in cadence and gear ratio. We chose cycling because its physics are well understood. In our physics-based model, we optimised for the only unknown, the drag number. For the neural network model, we used a simple recurrent neural network with one long-short-term-memory layer consisting of eight nodes. We found that the neural network model had a similar performance to that of the physics-based model.

Neural networks can help develop models when we do not understand the underlying principles of a problem. To develop the physics-based model we needed an understanding of the underlying principles of cycling. In comparison, the neural network could automatically learn the input-output relationship. If we would decide to add new input variables to the model, like for example continuous wind speed and direction, we suspect it would be easier for a scientist without an understanding of the underlying principles to train a new neural network with the added variables, than to find the correct way to add these variables to the physics-based model. Furthermore, a scientist with sufficient expertise in neural network modelling is well poised to develop new models for many other physical phenomena, without being an expert in the underlying principles of any of the systems. These are advantages of modelling using neural networks over physics-based modelling.

The physics-based model had more physically-meaningful parameters, introducing both advantages and disadvantages. For the physics-based model we used measured parameters, such as the rear wheel radius and the participant’s weight. We did not measure these parameters for the neural network, removing an extra step and potential for inaccuracies. But having fixed parameters makes it easier for future changes in the experimental environment. For example, if we decided to change the bike, the physics-based model would only need the new rear wheel radius and weight of the new bike, but the neural network would need new training with the new information. Having fixed physical parameters in the physics-based model also makes it more interpretable, which can help better understand the performance of a model, and subsequently increase the fundamental understanding of the particular problem itself. A neural network creates its own representation of a problem, which makes it harder to interpret.

Others have also shown the utility of neural networks when developing models of complex dynamic systems. Some have shown how to combine physics-based models with neural network models to enhance performance (Brunet et al., 2019; De Groote et al., 2022; Subraveti et al., 2022; Sun & Shi, 2022). Others have directly compared physics-based models and neural network models, similar to our project. For example, Choi et al. (Choi et al., 2020) compared the design and implementation of a physics-based model and neural network models for predicting the performance of a cooling system of a gasoline vehicle equipped with an electric control valve. Hu et al. (Hu et al., 2019) compared the design and implementation of a physics-based model, a combination of physics-based and neural network model, and a direct neural network model for simulating different metrics in a diesel combustion engine. In both cases, their findings suggest that developing the neural network required less expert knowledge and took significantly less time. In a complex process like this, expert knowledge is not only required to develop the physics-based model, but also to get valid measurements for the model’s parameters. Correspondingly, the authors found highest prediction accuracy with the neural network model. Such as in our project, they also see advantages in using physics-based models, such as better interpretability and better durability against the worst anomalous conditions in which there is not enough data for the neural network to learn accurately.

In our study, both models converged to similar, but not perfect, prediction accuracies. We expected this imperfect performance as there are many variables in cycling that affect the power output that we did not include in our models as they would have required complex measurement systems (Martin et al., 1998). For example, we did not measure or model headwinds or tailwinds which increase and decrease the drag force, respectively. Our participants cycled on an oval running track creating situations where in the presence of a prevailing wind, participants alternatively experienced headwinds and tailwinds with neither of these forces represented by changes in the inputs of the model. Incorporating continuous wind speed and direction could enhance the performance of both models. Doing so would be more difficult for the physics-based model, since we would have to understand how the wind speed and direction affects the whole system to properly add it to the model. In comparison, to add additional variables to the neural network model simply requires retraining the model with the added new variable in the input data. More generally, we suspect that as the complexity of the process to be modelled increases, or as the number of required measured inputs increases, a data-driven modelling approach will prove simpler, although harder to interpret, than a physical modelling approach.

## Supplemental Online Material

N/A

